# Imputation-Based Genomic Coverage Assessments of Current Genotyping Arrays: *Illumina HumanCore, OmniExpress, Multi-Ethnic global array and sub-arrays, Global Screening Array, Omni2.5M, Omni5M, and Affymetrix UK Biobank*

**DOI:** 10.1101/150219

**Authors:** Sarah C. Nelson, Jane M. Romm, Kimberly F. Doheny, Elizabeth W. Pugh, Cathy C. Laurie

**Affiliations:** Genetic Analysis Center, Dept. of Biostatistics, University of Washington; Center for Inherited Disease Research, Johns Hopkins University School of Medicine

## Abstract

Genotyping arrays have been widely adopted as an efficient means to interrogate variation across the human genome. Genetic variants may be observed either directly, via genotyping, or indirectly, through linkage disequilibrium with a genotyped variant. The total proportion of genomic variation captured by an array, either directly or indirectly, is referred to as “genomic coverage.” Here we use genotype imputation and Phase 3 of the 1000 Genomes Project to assess genomic coverage of several modern genotyping arrays. We find that in general, coverage increases with increasing array density. However, arrays designed to cover specific populations may yield better coverage in those populations compared to denser arrays not tailored to the given population. Ultimately, array choice involves trade-offs between cost, density, and coverage, and our work helps inform investigators weighing these choices and trade-offs.

## I. Introduction

Genotyping arrays have been widely adopted as an efficient means to interrogate variation across the human genome. Genetic variants may be observed either directly, via genotyping, or indirectly, through linkage disequilibrium (LD) with a genotyped variant. The total proportion of genomic variation captured by an array, either directly or indirectly, is referred to as “genomic coverage.” Assessments of genomic coverage generally take one of two forms. (1) LD-based genomic coverage estimates describe the fraction of variants that are in LD with array variants at a given pairwise correlation value (r^2^ ≥ 0.8, e.g.)^1,2^. (2) Imputation-based genomic coverage leverages array variants to impute into a more densely genotyped or sequenced reference panel, such as the HapMap Project^3^ or 1000 Genomes Project^4,5^. In this approach, genomic coverage is quantified as the proportion of variants with an imputation r^2^ above a given threshold (again, usually r^2^ ≥ 0.8)^6^.

In response to both the evolving needs of the genetic research community and technological advances in genotyping, new arrays from both Illumina (www.illumina.com) and Affymetrix (www.affymetrix.com) continue to be developed. While vendors often provide genomic coverage estimates in product documentation, the methods may not be comparable and/or well-described, making it difficult to objectively compare across arrays. Previously we published imputation-based genomic coverage for a series of arrays using Phase 1 of the 1000 Genomes Project as a reference^7^. Here we report an extension of those previous coverage analyses, using 1000 Genomes Phase 3^5^ and an updated set of arrays, including the Illumina Multi-Ethnic arrays and Global Screening Array. The original and present analyses are intended to function as practical guides to researchers planning genetic studies. Both coverage and accuracy information are presented.

Product literature from the manufacturers details the design and intended uses of the arrays we assess here. In brief, Illumina Multi-Ethnic arrays were designed using content from multiple sources, including Phase 3 of the 1000 Genomes Project, the Consortium on Asthma among African ancestry Populations in the Americas (CAAPA), and Population Architecture using Genomes and Epidemiology (PAGE). The Multi-Ethnic global array is intended for use in studies with diverse ancestry, while the sub-arrays have genome-wide tagging content tailored to certain populations. Specifically, the AFR/AMR sub-array is optimized for African American and Hispanic study samples, while the EUR/EAS/SAS sub-array is optimized for European, East Asian, and South Asian samples. For more details, see the array-specific data sheets available at http://www.illumina.com/products/infinium-multi-ethnic-global-array.html. The Illumina Global Screening Array (GSA) was developed by a consortium of human genetics researchers to function as a cost-efficient GWAS array with additional clinically-relevant content (see https://www.illumina.com/content/dam/illumina-marketing/documents/products/datasheets/infinium-commercial-gsa-data-sheet-370-2016-016.pdf). The Affymetrix UK Biobank array is optimized for “populations of European and British ancestry” and indeed was initially designed for use in the UK Biobank (see http://www.affymetrix.com/support/technical/datasheets/uk_axiom_biobank_genotyping_arrys_datasheet.pdf). The Illumina HumanCore, OmniExpress, Omni2.5M and Omni5M are included as earlier-generation arrays to serve as reference points for coverage.

## II. Materials and Methods

For these genomic coverage assessments, we used the publicly available 1000 Genomes Project Phase 3 integrated variant set, in variant call format (VCF, available at http://ftp.1000genomes.ebi.ac.uk/vol1/ftp/release/20130502/). Samples are grouped into continental panels or “super-populations”^5^: African (AFR), Americas (AMR), East Asian (EAS), European (EUR), and South Asian (SAS) — see Table 1. Following our published methods^7^, we first randomly assigned each of the 2,504 Phase 3 samples into one of ten batches, balancing samples across populations. We then created subsets of the data in the 1000 Genomes VCF files, which are separated by chromosome. For each sample batch, we extracted genotypes from the VCF files for just the samples in the batch and the variants on the array. Array variants were identified within the VCF using chromosome and base pair positions from the most current array manifests available from vendor websites (see Table 2). A graphical overview of our methods is shown in Figure 1.

**Figure 1.**
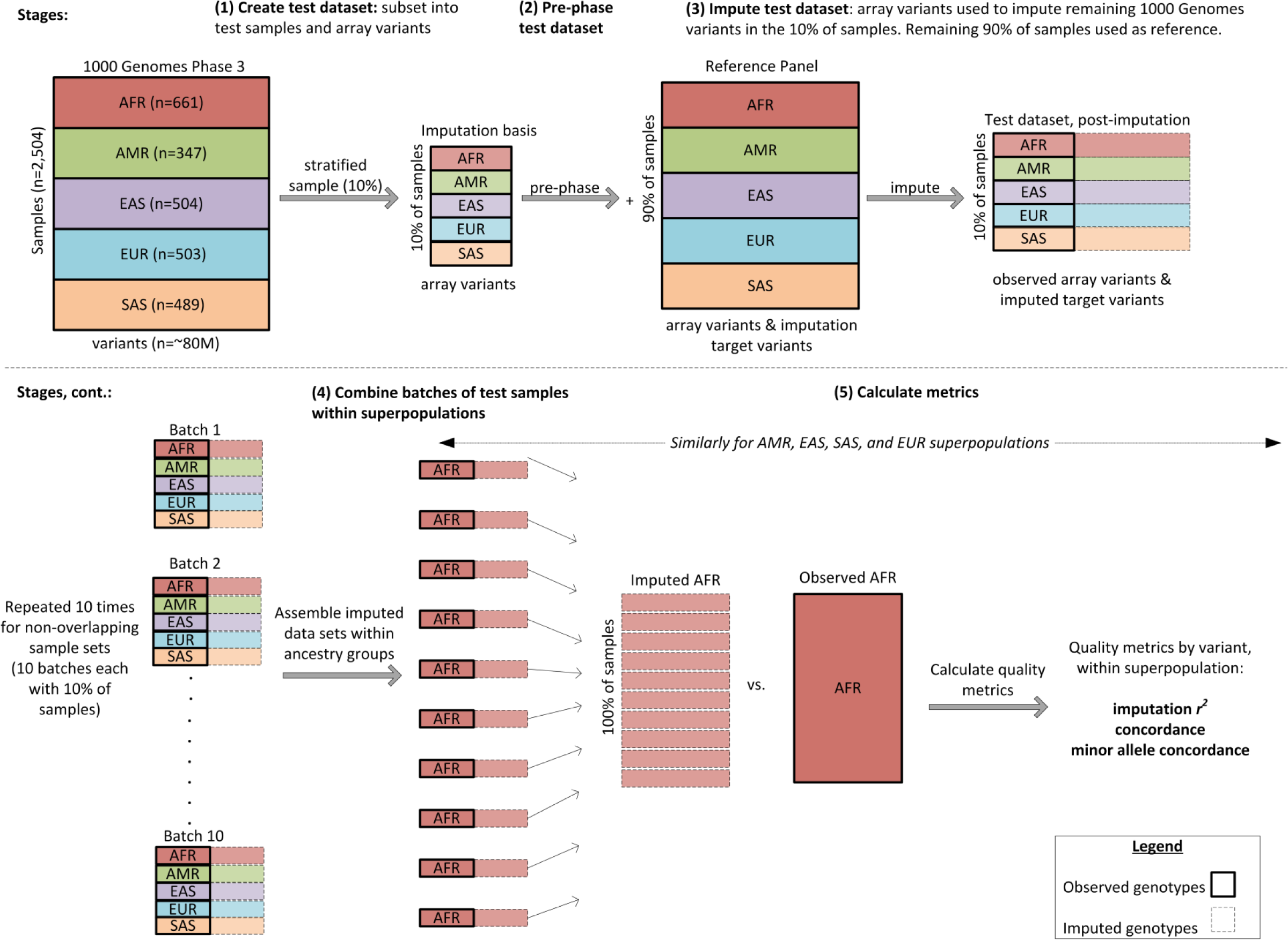
Schematic of the study design used to assess genomic coverage of select genotyping arrays, with 1000 Genomes Project Phase 3 as a source of both test and reference data. Dashed lines indicate imputed genotypes; solid lines indicate observed genotypes. Adapted from Figure 1 in Nelson et al.^7^.

**Table 1.** An overview of the 2,504 samples in the 1000 Genomes Project Phase 3 reference used for these coverage assessments. The Project assigned each population to one of five super-populations: African (AFR), Americas (AMR), East Asian (EAS), European (EUR), and South Asian (SAS). Each sample was assigned to one of ten test batches at random, yet keeping an even representation of populations across batches.

| Full Population Name | Abbreviation | Number of Samples | Number of samples per batch* |
| --- | --- | --- | --- |
| African Caribbean in Barbados | ACB | 96 | 10 |
| African Ancestry in Southwest US | ASW | 61 | 6 |
| Esan in Nigeria | ESN | 99 | 10 |
| Gambian in Western Division, The Gambia | GWD | 113 | 11 |
| Luhya in Webuye, Kenya | LWK | 99 | 10 |
| Mende in Sierra Leone | MSL | 85 | 8 |
| Yoruba in Ibadan, Nigeria | YRI | 108 | 11 |
| *Total African ancestry* | *AFR* | *661* | *66* |
| Colombian in Medellin, Colombia | CLM | 94 | 9 |
| Mexican ancestry in Los Angeles, California | MXL | 64 | 6 |
| Peruvian in Lima, Peru | PEL | 85 | 8 |
| Puerto Rican in Puerto Rico | PUR | 104 | 10 |
| *Total Americas ancestry* | *AMR* | *347* | *33* |
| Chinese Dai in Xishuangbanna, China | CDX | 93 | 9 |
| Han Chinese in Beijing, China | CHB | 103 | 10 |
| Southern Han Chinese, China | CHS | 105 | 10 |
| Japanese in Tokyo, Japan | JPT | 104 | 10 |
| Kinh in Ho Chi Minh City, Vietnam | KHV | 99 | 10 |
| *Total East Asian ancestry* | *EAS* | *504* | *49* |
| Bengali in Bangladesh | BEB | 86 | 9 |
| Gujarati Indian in Houston, TX | GIH | 103 | 10 |
| Indian Telugu in the UK | ITU | 102 | 10 |
| Punjabi in Lahore, Pakistan | PJL | 96 | 10 |
| Sri Lankan Tamil in the UK | STU | 102 | 10 |
| *Total South Asian ancestry* | *SAS* | *489* | *49* |
| Utah residents with Northern and Western European ancestry | CEU | 99 | 10 |
| Finnish in Finland | FIN | 99 | 10 |
| British in England and Scotland | GBR | 91 | 9 |
| Iberian populations in Spain | IBS | 107 | 11 |
| Toscani in Italia | TSI | 107 | 11 |
| *Total European ancestry* | *EUR* | *503* | *51* |

\*Sample counts are for batches 1-9; batch 10 took the remainder, and thus each population contributed slightly more or fewer samples.

**Table 2.** Summaries of each array included in these genomic coverage analyses. “Product information” refers to the manifest version; “Date manufactured” and “Loci Count” were extracted from the array manifest headers. The last two columns indicate the percentage of unique positions assayed by the array (mapped to chromosomes 1 through 22 and the non-pseudoautosomal portion of the X chromosome) that are also found in the 1000 Genomes Phase 3 (“1000G Ph3”) integrated variant set. The second to last column is over all 1000 Genomes variants while the last column is restricted to those with a minimum of two minor allele copies seen in at least one of the five super-populations. Because the last column is over a restricted set of 1000 Genomes variants, the percentage overlap decreases. Note coverage analyses were restricted to chromosomes 1 and 22. Arrays are manufactured by Illumina, with the exception of the Affymetrix UK Biobank.

| Array Name | Product information | Date Manufactured | Loci Count | % overlap w/ 1000G Ph3 | % overlap w/1000G Ph3, MAF filtered |
| --- | --- | --- | --- | --- | --- |
| HumanCore | 24v1-0_A | 21-Aug-14 | 306,670 | 99% | 99% |
| Global Screening Array | 24v1-0_A | 22-Oct-16 | 642,824 | 98% | 96% |
| OmniExpress | 24v1-1_A | 14-Oct-14 | 713,014 | 99% | 99% |
| Affymetrix UK Biobank | na 35 | 9-Mar-15 | 820,967 | 94% | 92% |
| Multi-ethnic AMR/AFR | 8v1-0_A1 | 7-Dec-15 | 1,430,141 | 85% | 81% |
| Multi-ethnic EUR/EAS/SAS | 8v1-0_A1 | 7-Dec-15 | 1,475,140 | 88% | 85% |
| Multi-ethnic global | _A1 | 20-Oct-15 | 1,779,819 | 86% | 83% |
| Omni2.5M | 8v1-2_A1 | 6-Mar-15 | 2,338,671 | 95% | 94% |
| Omni 5M | 4-v1-1_A | 11-Feb-15 | 4,284,426 | 96% | 95% |

We removed the VCF phasing information when converting to PLINK binary format (via VCFtools^8^) to better approximate array genotypes, which are usually at first unphased. We then used SHAPEIT2^9^ (v2.r790) software to pre-phase each dataset. The resulting phased haplotypes were then imputed with IMPUTE2^10^ (v2.3.2) where, for each batch, the remaining 1000 Genomes samples served as a worldwide imputation reference panel. We restricted imputation to variants with at least two copies of the minor allele in any one of the five super-populations. Following imputation, the output (genotype probabilities file) for each of the ten sample batches was then combined. We restricted these experiments to chromosomes 1 and 22.

To assess imputation accuracy and by extension genomic coverage, we compared imputed results at all the non-array variants to observed genotypes from the original VCF files. This comparison was performed separately by super-population and restricted to variants with at least two copies of the minor allele in the given panel, yielding 2,701,990 variants in the AFR panel; 1,709,647 in AMR; 1,304,742 in EAS; 1,401,215 in EUR; and 1,556,550 in SAS (note counts are across chromosomes 1 and 22). This variant restriction was done to avoid an excess of missing imputation metrics, which occur when a variant is either imputed or observed to be monomorphic. Imputation included all three variant types in the 1000 Genomes Project release: single nucleotide, insertion/deletion, and structural variants, and thus all three variant types were included in these metrics calculations.

For each variant we calculated three metrics: (1) the squared correlation between observed and imputed allelic dosage, which we call “dosage r^2^”; (2) the concordance between observed and most likely imputed genotype, the “genotype concordance”; and (3) the concordance between observed and most likely imputed genotype, when at least one of those two genotypes contains one or two copies of the minor allele, which we call “minor allele (MA) concordance.” All metrics were calculated using R statistical and graphing software^11^. Array (observed) variants are included in these metrics summaries and are given dosage r^2^, genotype concordance, and MA concordance values of 1.

## III. Results and Discussion

Figure 2 shows the fraction of variants passing a dosage r^2^ threshold of 0.8. Each quadrant represents a different minor allele frequency (MAF) bin, where MAF was calculated in each super-population. Genomic coverage assessments commonly use the MAF groupings of >1% and >5%, which we have shown in panels C and D, respectively. The position of arrays along the x-axis indicates the total number of unique positions assayed by each array. Thus the least dense array is to the far left (Illumina HumanCore) and the densest array to the far right (Illumina Omni5M). Figure 3 shows mean MA concordance, by MAF bin and super-population. Similarly Figure 4 shows mean dosage r^2^, and Figure 5 shows genotype concordance. All the data plotted in Figures 2 – 5 are presented numerically in Tables 3A-E, which also include tabular summaries of mean dosage r^2^ and genotype concordance. Tables 3A-E also indicate the counts of variants in each MAF bin, as this count differs by super-population. The same metrics are also presented in barplots in Appendix A, for an alternative visualization.

**Figure 2.**
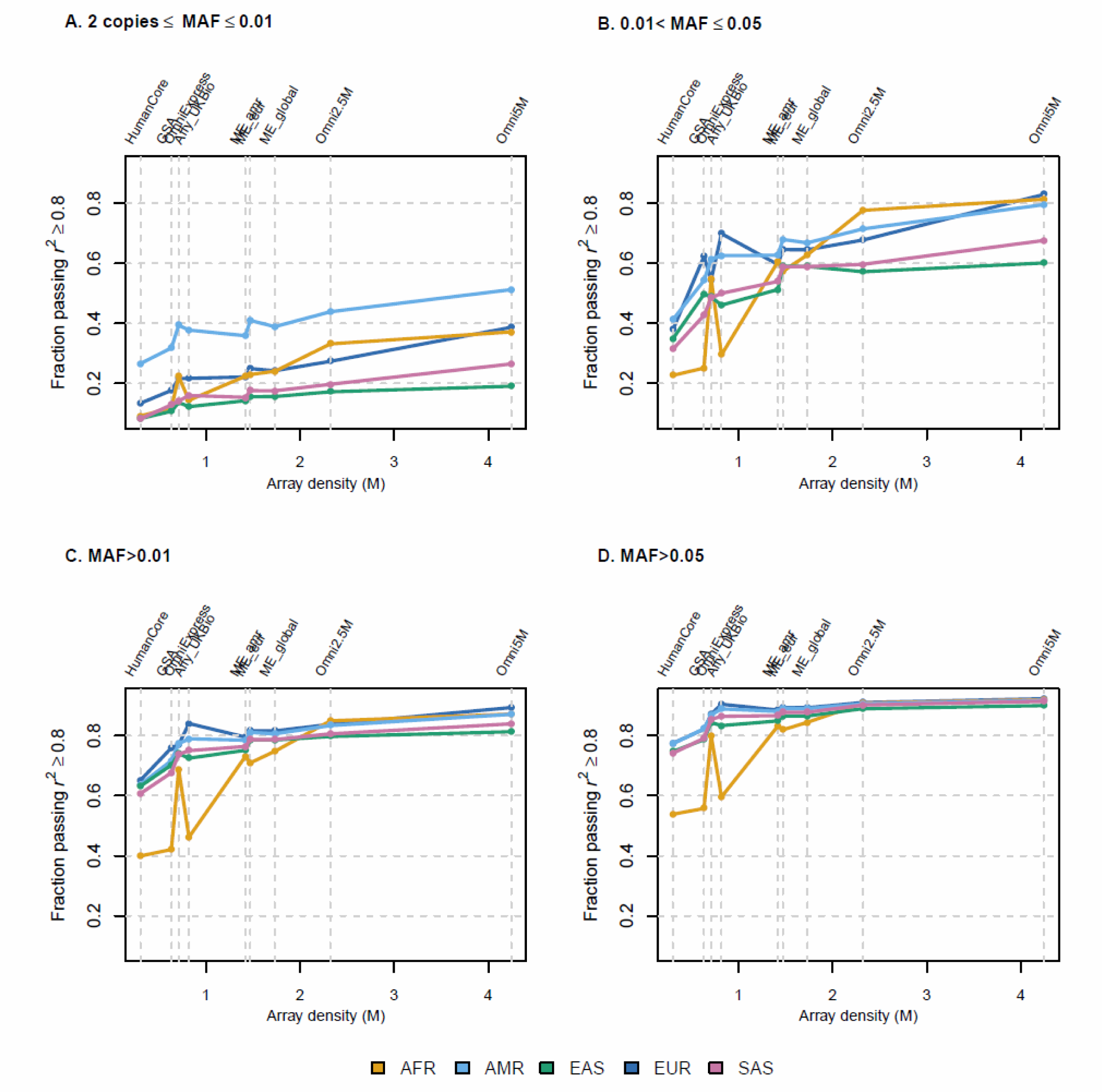
Fraction of variants passing a dosage r^2^ threshold of 0.8, by MAF bin and super-population. Each of the five super-populations, indicated by a different colored line, was divided into ten batches and imputed using all remaining samples as a worldwide reference panel. The dosage r^2^ metric plotted here is the squared correlation between imputed and observed allelic dosage in the samples comprising the super-population. The y-axis is the proportion of variants (imputed and observed) with dosage r^2^ ≥ 0.8, restricted to variants with at least two copies of the minor allele in the given super-population. The x-axis position of each array corresponds to the number of unique positions assayed by that array. Thus the order of the arrays on each axis is as follows: HumanCore, Global Screening Array, OmniExpress, Affymetrix UK Biobank, Multi-Ethnic AMR/AFR, Multi-Ethnic EUR/EAS/SAS, Multi-Ethnic global, Omni2.5M, and Omni5M.

**Figure 3.**
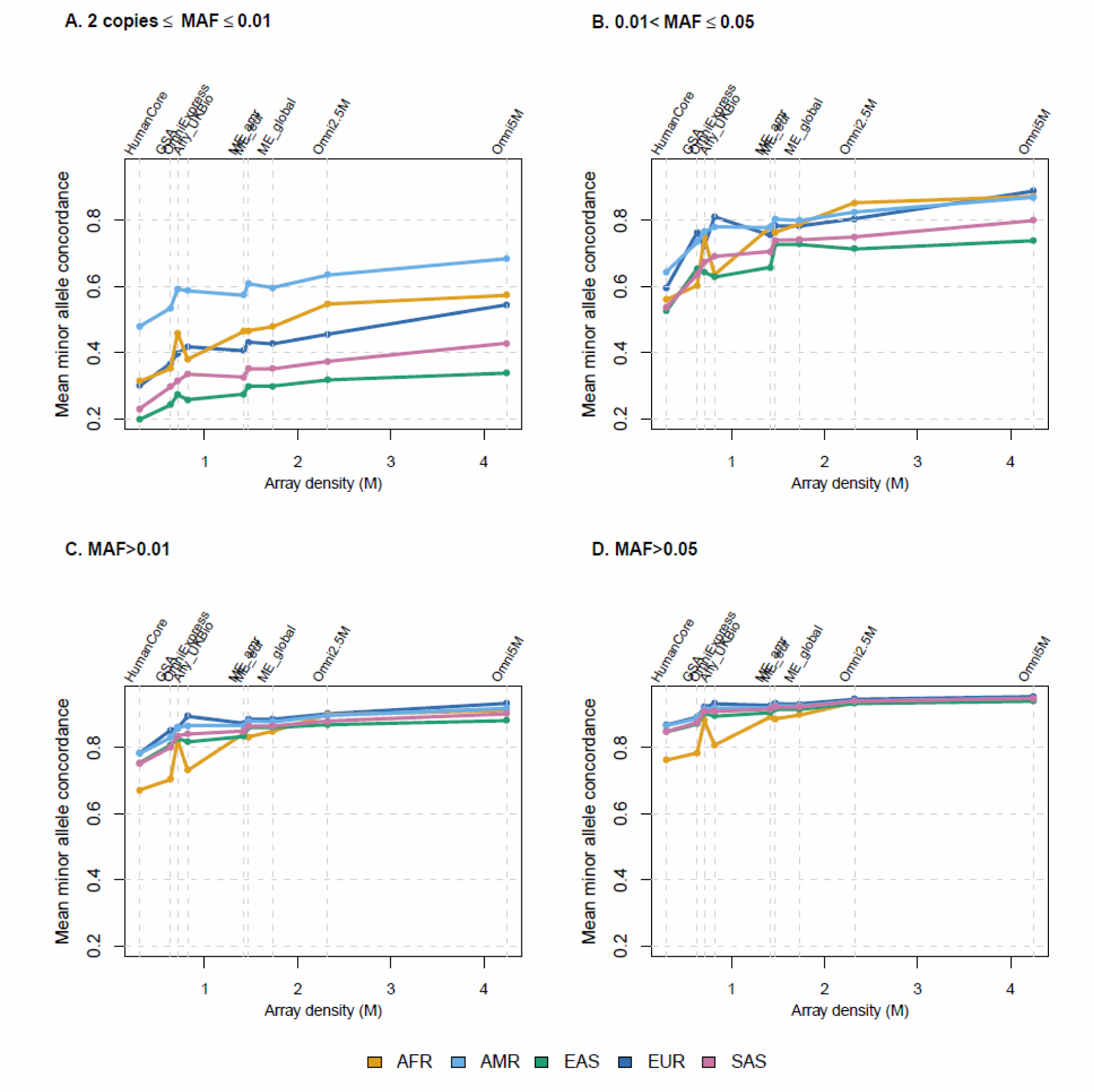
Mean minor allele (MA) concordance, by MAF bin and super-population. Each of the five super-populations, indicated by a different colored line, was divided into ten batches and imputed using all remaining samples as a worldwide reference panel. The y-axis values are mean MA concordance in samples comprising the given super-population. MA concordance is defined as the concordance (percent agreement) between observed and most likely imputed genotype, when at least one of those two genotypes contains one or two copies of the minor allele. Variants were restricted to those with at least two copies of the minor allele in the given super-population. Both imputed and observed variants are included in this average; the latter with MA concordance values of 1. The x-axis position of each array corresponds to the number of unique positions assayed by that array. Thus the order of the arrays on each axis is as follows: HumanCore, Global Screening Array, OmniExpress, Affymetrix UK Biobank, Multi-Ethnic AMR/AFR, Multi-Ethnic EUR/EAS/SAS, Multi-Ethnic global, Omni2.5M, and Omni5M.

**Figure 4.**
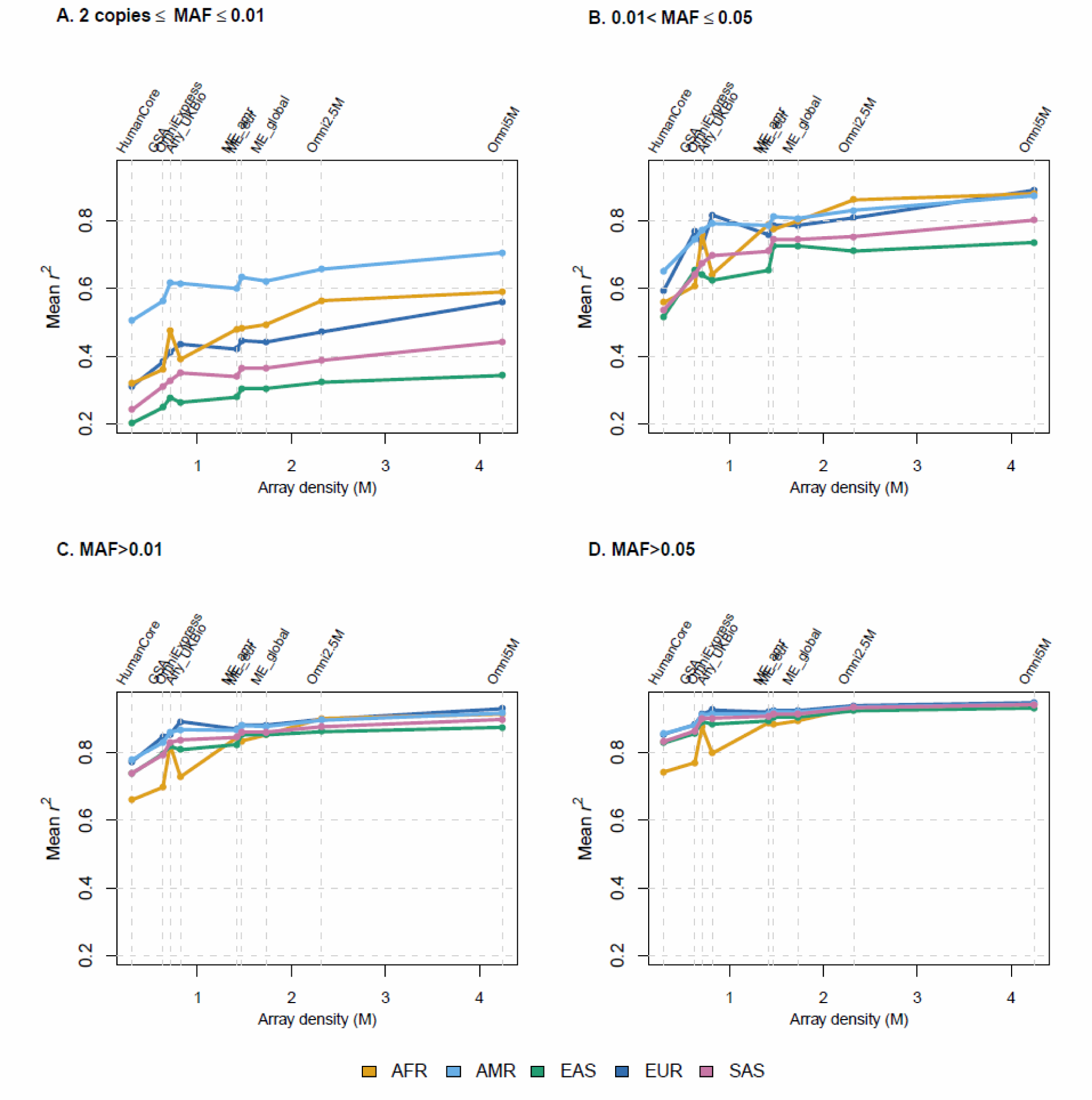
Mean dosage r^2^, by MAF bin and super-population. Each of the five super-populations, indicated by a different colored line, was divided into ten batches and imputed using all remaining samples as a worldwide reference panel. The y-axis is the mean dosage r^2^ metric: the squared correlation between imputed and observed allelic dosage in the samples comprising the super-population. The x-axis position of each array corresponds to the number of unique positions assayed by that array. Thus the order of the arrays on each axis is as follows: HumanCore, Global Screening Array, OmniExpress, Affymetrix UK Biobank, Multi-Ethnic AMR/AFR, Multi-Ethnic EUR/EAS/SAS, Multi-Ethnic global, Omni2.5M, and Omni5M.

**Figure 5.**
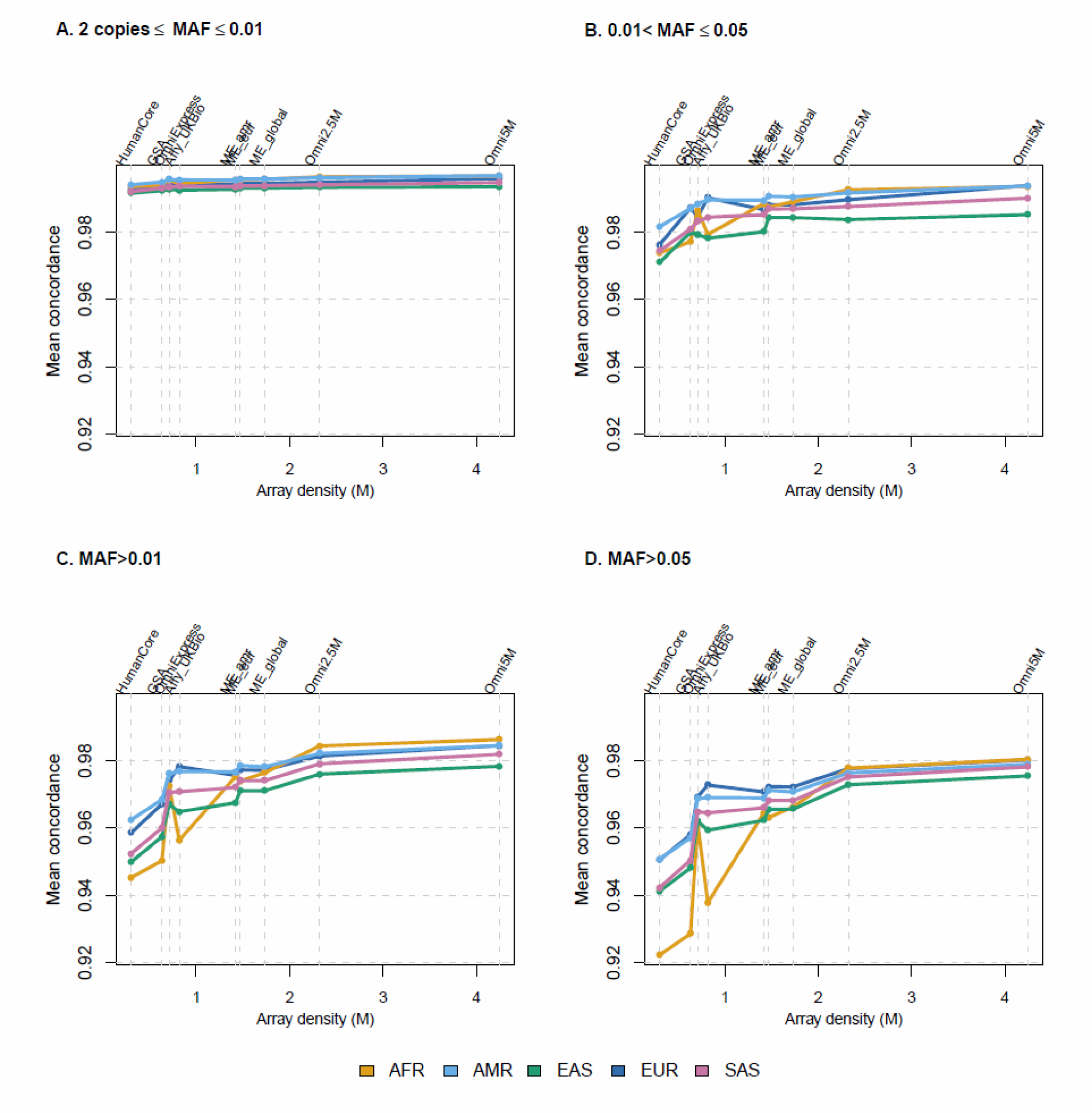
Mean genotype concordance, by MAF bin and super-population. Each of the five super-populations, indicated by a different colored line, was divided into ten batches and imputed using all remaining samples as a worldwide reference panel. The y-axis values are mean genotype concordance in samples comprising the given super-population. Variants were restricted to those with at least two copies of the minor allele in the given super-population. Both imputed and observed variants are included in this average; the latter with concordance values of 1. The x-axis position of each array corresponds to the number of unique positions assayed by that array. Thus the order of the arrays on each axis is as follows: HumanCore, Global Screening Array, OmniExpress, Affymetrix UK Biobank, Multi-Ethnic AMR/AFR, Multi-Ethnic EUR/EAS/SAS, Multi-Ethnic global, Omni2.5M, and Omni5M.

**Tables 3A-3E.**
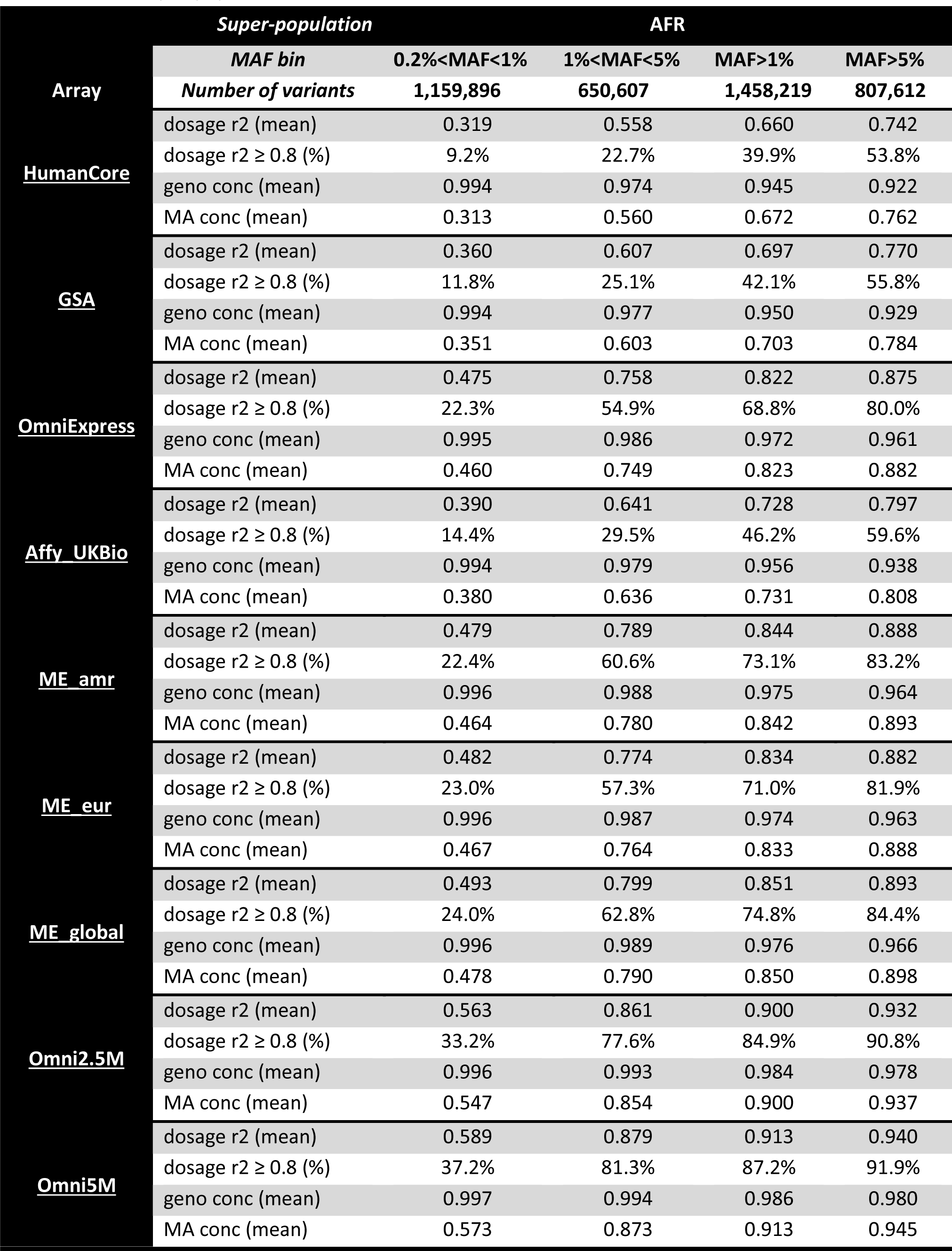
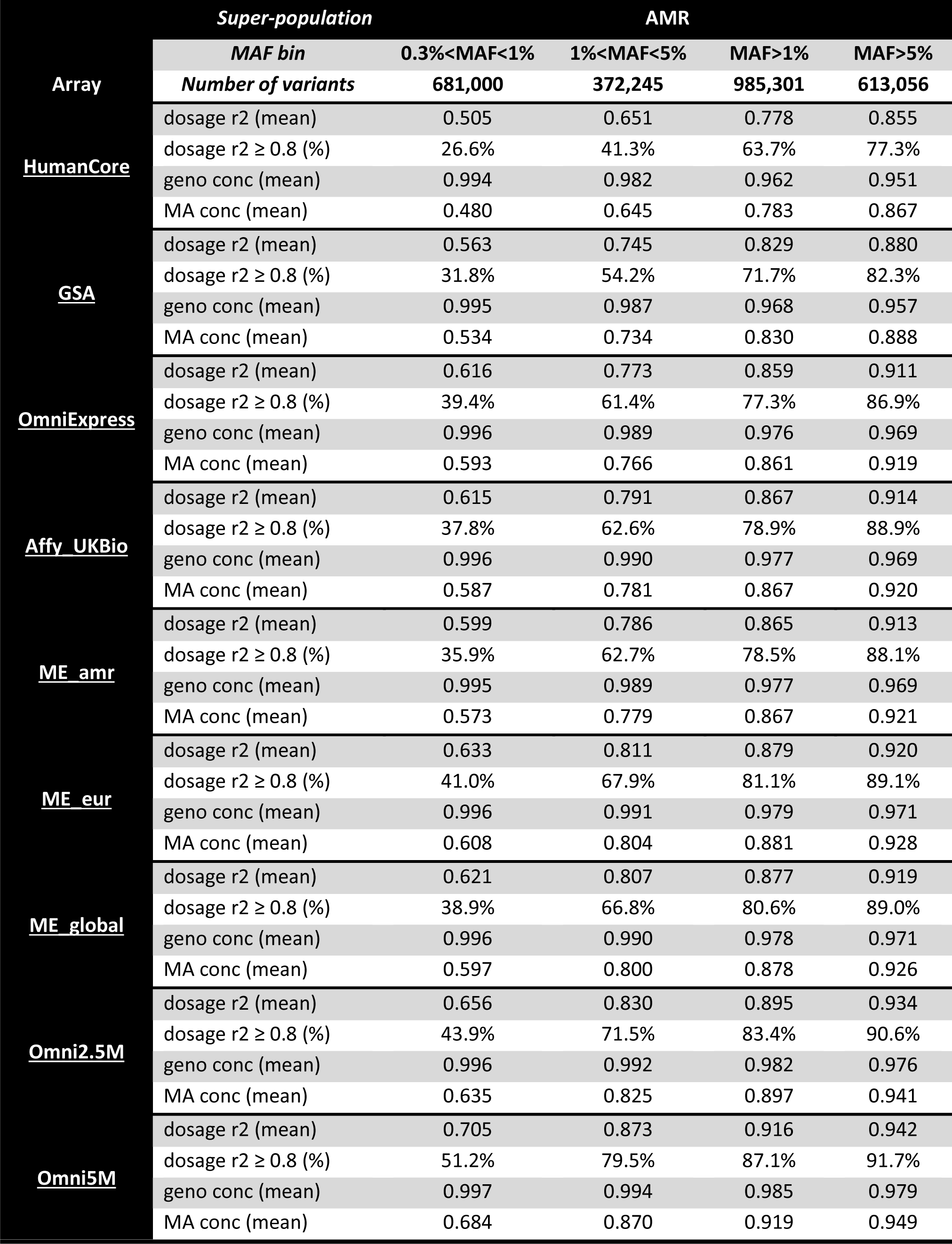
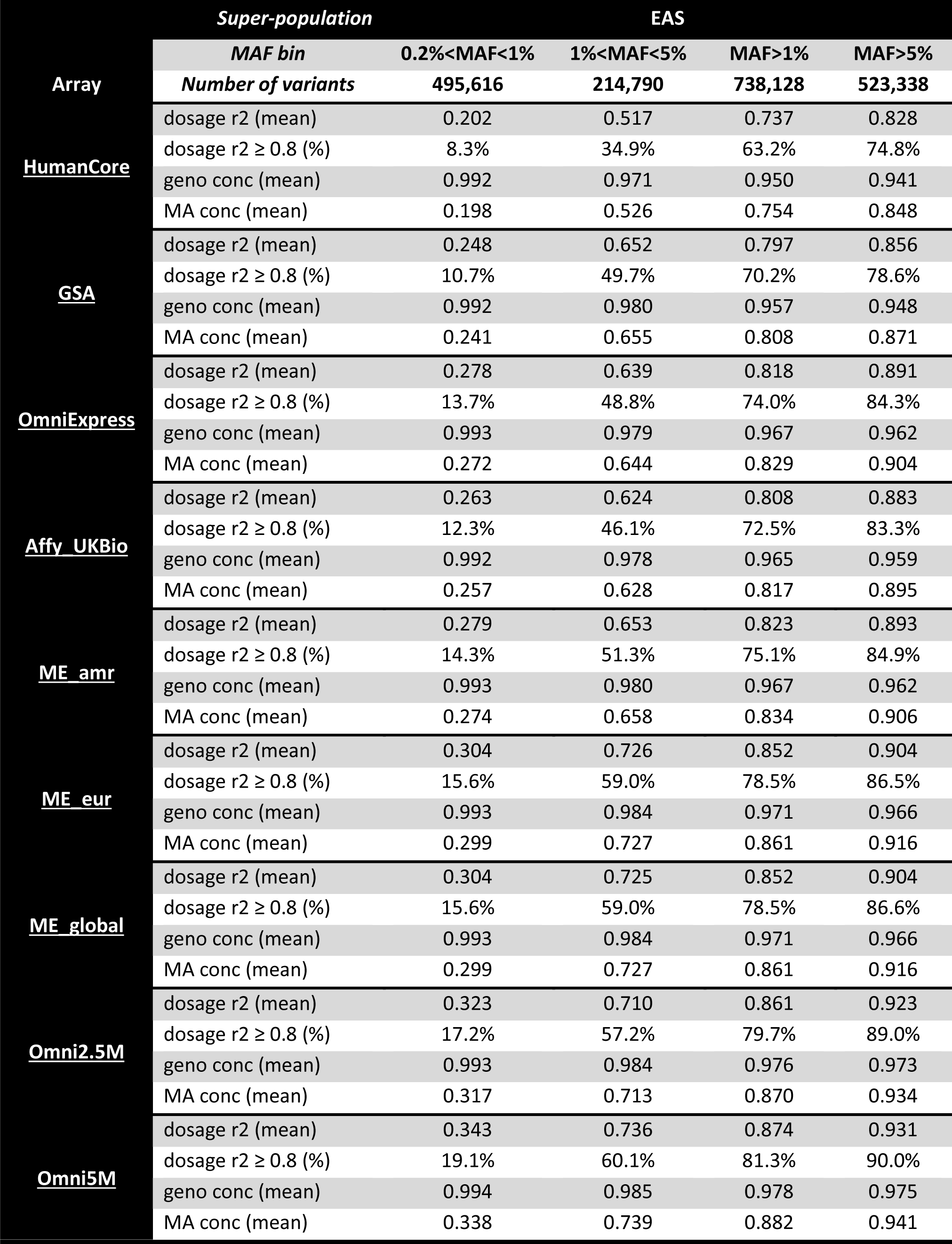
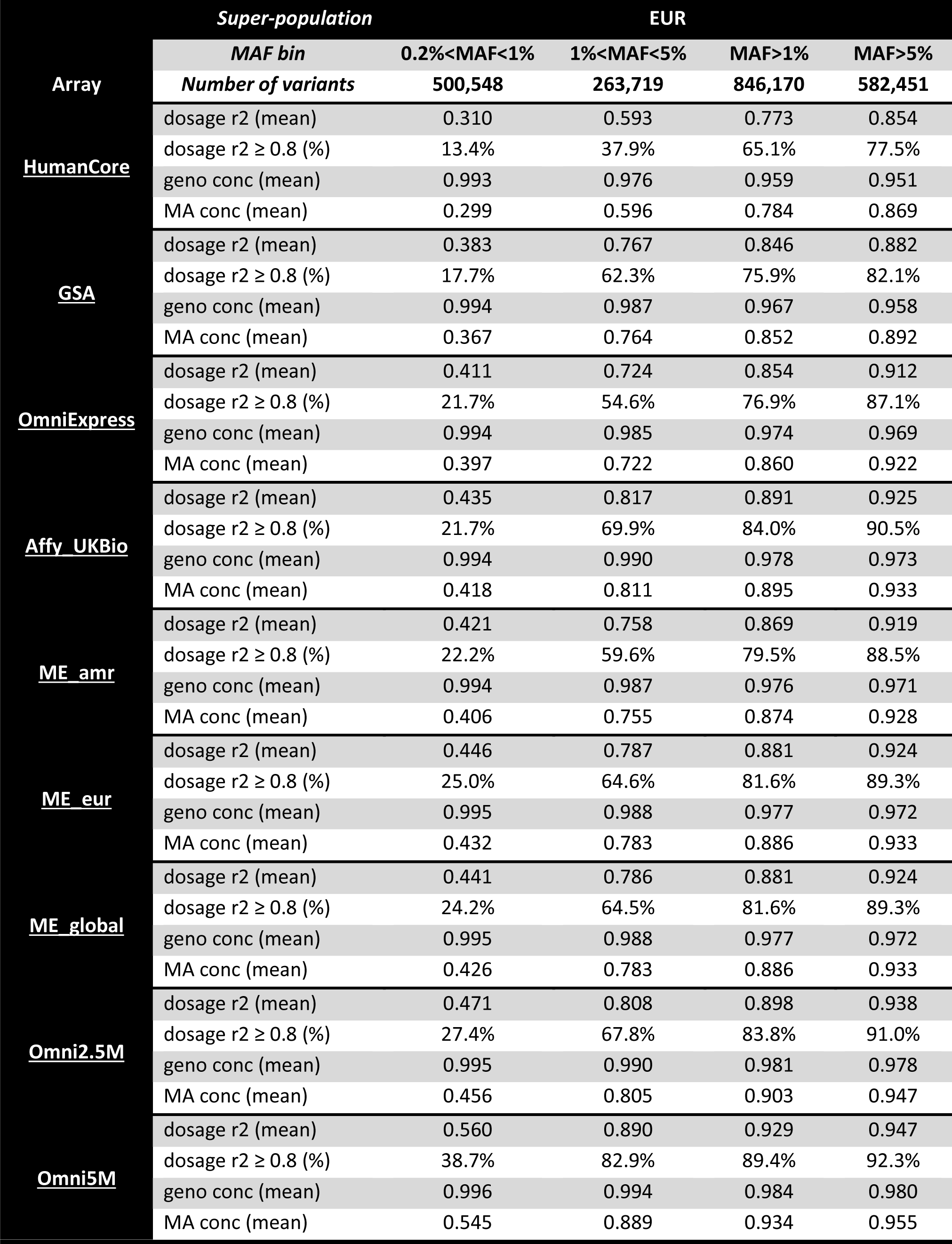
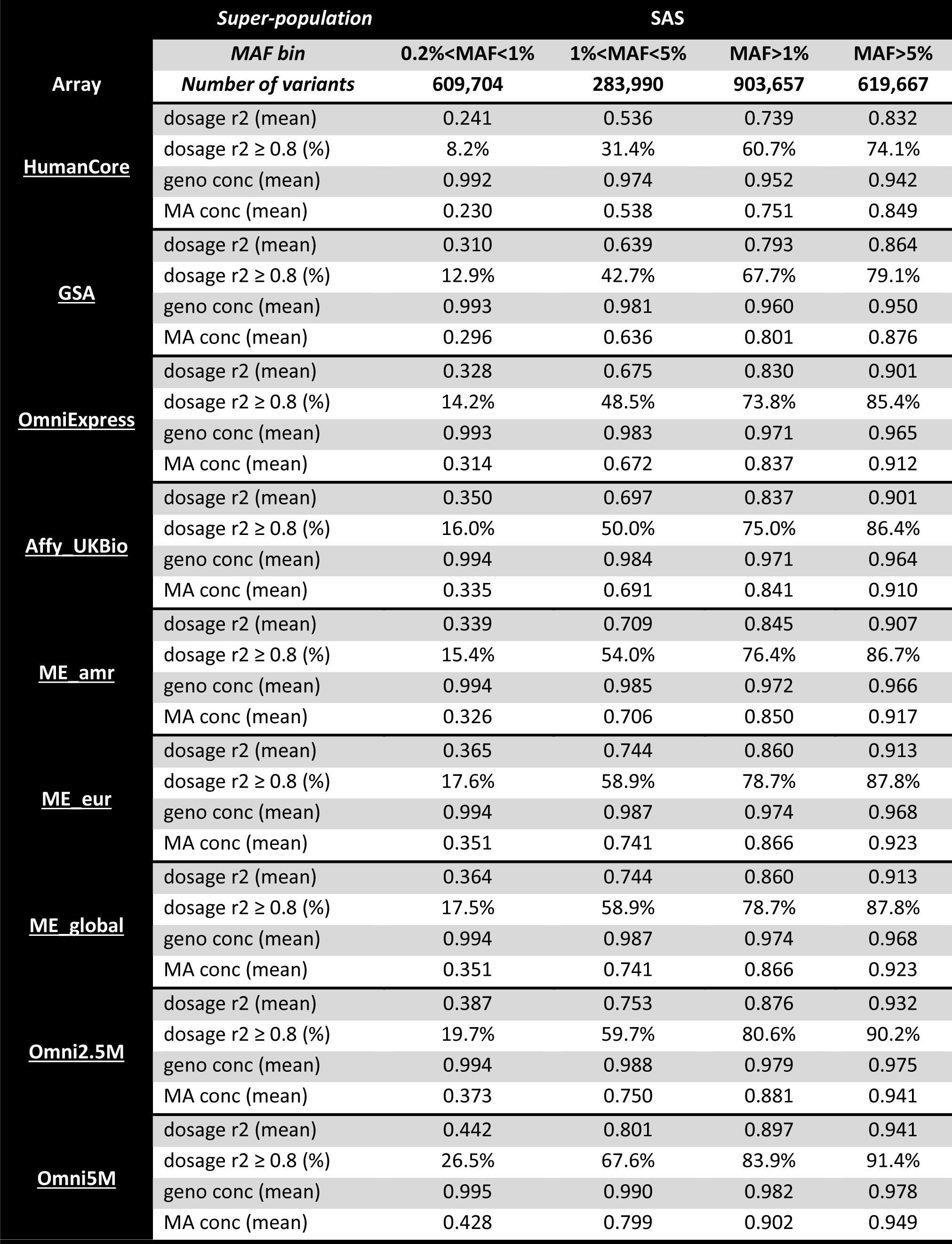
Genome-wide coverage estimates for all evaluated arrays, separately by MAF bin and super-population. “Number of variants” gives the count of 1000 Genomes Phase 3 variants per MAF bin for the given super-population. This count includes “observed” (i.e., included on the assessed array) variants, which have dosage *r^2^* and concordance values of 1. The “geno conc” is the genotype concordance; “MA conc” is the minor allele concordance (see narrative for details). **Table 3A.** AFR super-population **Table 3B.** AMR super-population **Table 3C.** EAS super-population **Table 3D.** EUR super-population **Table 3E.** SAS super-population

As one might expect, quality metrics generally improve with increasing array density, regardless of either super-population or MAF bin. Rare variants (MAF < 1%) are generally not well imputed by either low or high density arrays. Common (MAF> 5%) variants are generally well imputed by all arrays, though there is still a distinction between the sparsest and densest arrays. Most of the line plots are monotonic. Some notable exceptions are the transitions between (1) OmniExpress and Affy UK Biobank in the AFR super-population and between (2) Affy UK Biobank and the Multi-Ethnic arrays in the EUR super-population.

The Multi-Ethnic global array and sub-arrays each assay a similar number of variants, though the global array has a slightly higher density. According to product literature, the Multi-Ethnic AMR/AFR array (“ME_amr” in plots) is designed for use in Hispanic and African American populations, while the Multi-Ethnic EUR/EAS/SAS array (“ME_eur” in plots) is designed for use in European, East Asian, and South Asian populations. Notably, according to these experiments, the Multi-Ethnic AMR/AFR array does not consistently outperform Multi-Ethnic EUR/EAS/SAS array for the AFR and AMR 1000 Genomes panels. However, this may be because populations in AMR and AFR are imperfect proxies for contemporary Hispanic and African American populations in the US. In contrast, for EUR, EAS, and SAS panels, the Multi-Ethnic EUR/EAS/SAS does perform slightly better than Multi-Ethnic AMR/AFR, which is consistent with the stated purpose of the arrays. While denser, the Multi-Ethnic global array does not consistently outperform the subarrays, though the relative performances vary by super-population and MAF bin.

The GSA results are generally consistent with what one would expect based on array densities: coverage is better than the less dense HumanCore but not as good as for the OmniExpress, the next densest assessed array after the GSA. One situation in which GSA outperforms the OmniExpress is for MAF 1-5% variants in the EUR and, to a lesser extent, the EAS super-populations. Notably, in contrast to the mostly tagging-focused OmniExpress, the GSA devotes ~10% of its content to clinically-relevant markers. These clinically-relevant markers may be underperforming as an imputation basis but derive value from their clinical associations. There are several potential limitations to these analyses. First, by assessing genomic coverage in the context of genome-wide imputation, we can only assess array variants that are also present in the chosen reference panel (1000 Genomes Phase 3 integrated variant set). Thus not all array variants can inform the imputation, for one of two reasons: (1) the array variant is not present in this 1000 Genomes Project dataset or (2) the array variant is present but without two or more copies of the minor allele in any one of the five super-populations (the MAF filtering threshold for imputation). In Table 2 we show for each array the percent overlap with 1000 Genomes at any allele frequency and the percent overlap with the 1000 Genomes with the requisite minor allele count to be included in imputation. Notably, the Multi-Ethnic arrays have less overlap than the other arrays.

A second caveat is that we used chromosomes 1 and 22 to estimate genome-wide coverage. Above we present coverage for the two chromosomes combined. To assess whether coverage patterns were consistent across these two chromosomes, we also evaluated them separately. We observed a statistically significant different between the two chromosomes for all four coverage metrics. However, the trends in coverage for the two chromosomes are quite consistent across arrays, for each of the super-populations and MAF bins. These patterns are illustrated in Figure 6. While the absolute coverage metrics may differ, the relative coverage is similar — evidence that supports the reliability of our approach here. Furthermore, it is not surprising that coverage may differ by chromosome, given that chromosomes are heterogeneous with respect to structure, gene density, and CG content^12^. This heterogeneity of features may lead to differential destiny of genotyping assays on a SNP array and thus differences in coverage across chromosomes. Other factors, such as LD patterns, may also vary among chromosomes. Our primary goal here is to get a sense of how arrays compare to each other and to observe the trade-offs in assay density and genomic coverage. (Note GSA is absent from these chromosome-specific analyses, due to timing of coverage experiments. However, given the consistency of the chromosome-specific trends across the remaining arrays, it is reasonable to expect the same trends would hold for GSA.)

**Figure 6.**
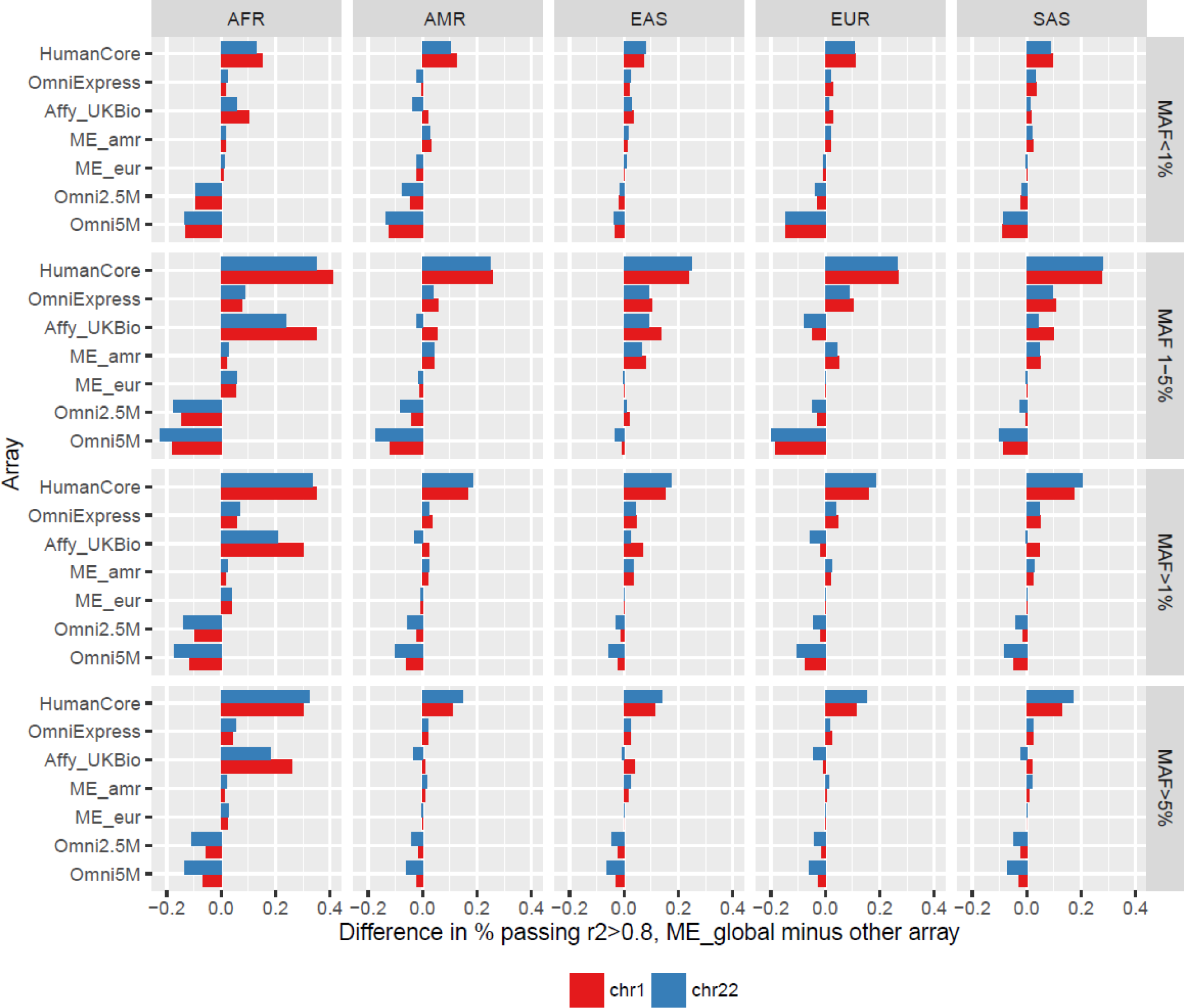
Comparison of genomic coverage on chromosomes 1 and 22. Each plot in the grid is for a given MAF group (as rows) and super-population (as columns). The bars show the difference in fraction of variants with r^2^> 0.8, subtracting the value for the array indicated from the Multi-Ethnic global array. Note the Multi-Ethnic global array is omitted from the plot, as the comparison is always 0% difference. Positive values on the x-axis mean Multi-Ethnic global has higher coverage; negative values means the labelled array has higher coverage. The Global Screening Array was not assessed for chromosome-specific metrics and is thus not represented on this plot.

In summary, our current and previous^7^ imputation-based genomic coverage analyses are intended to help researchers weigh the costs and benefits of different array choices across different genetic ancestry groups. Ultimately, the genomic coverage afforded by a given array will likely be affected by many study-specific factors, such as genotyping quality, genetic ancestry profile of study samples, and imputation procedures. These findings, however, can serve as a robust starting point for researchers evaluating their genotyping array options.

## Appendix A.

Barplots of coverage metrics, by super-population. Arrays are ordered by density within each MAF group, with least dense to the left (HumanCore) and most dense to the right (Omni5M). The array color coding is as follows:

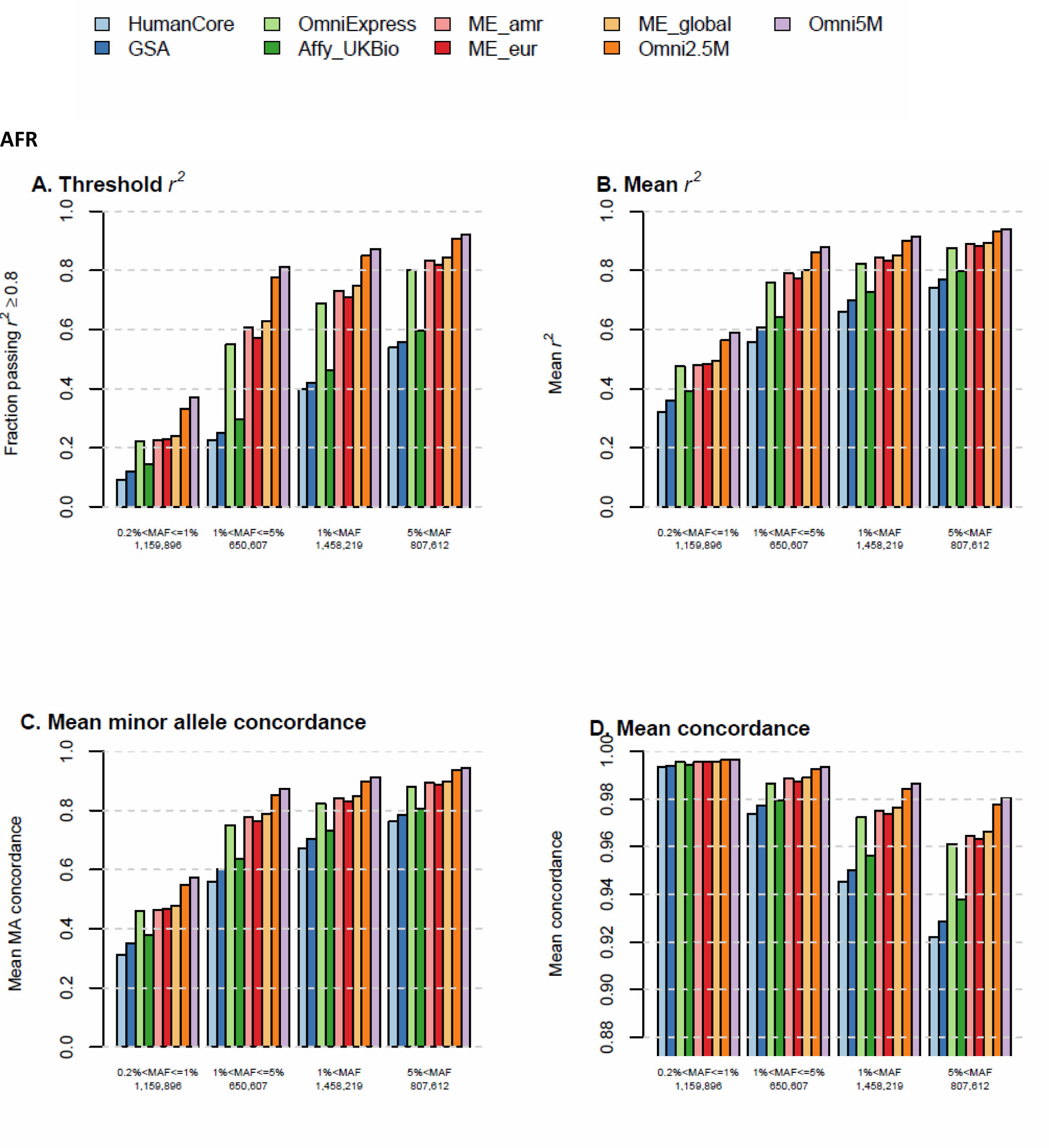

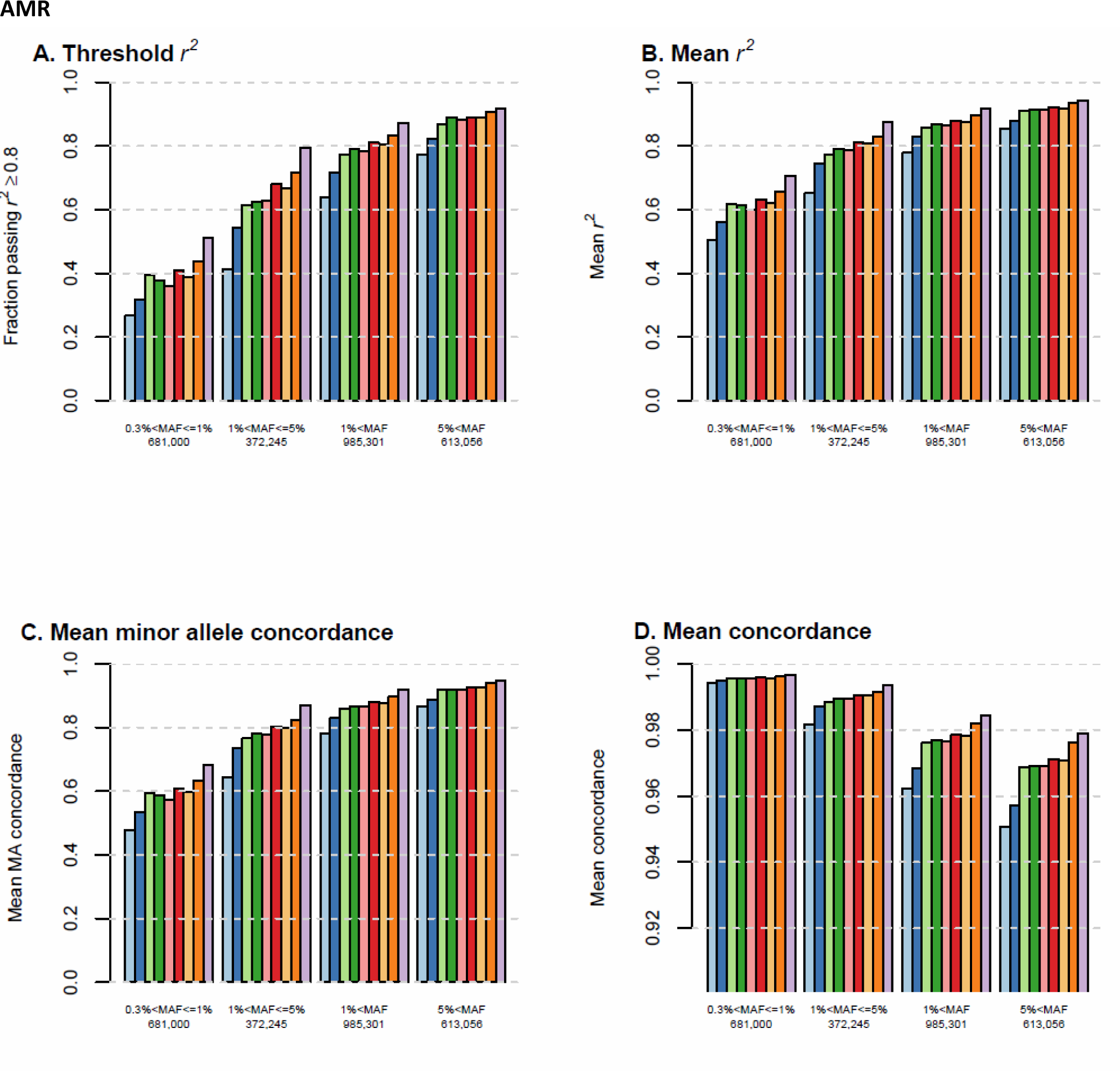

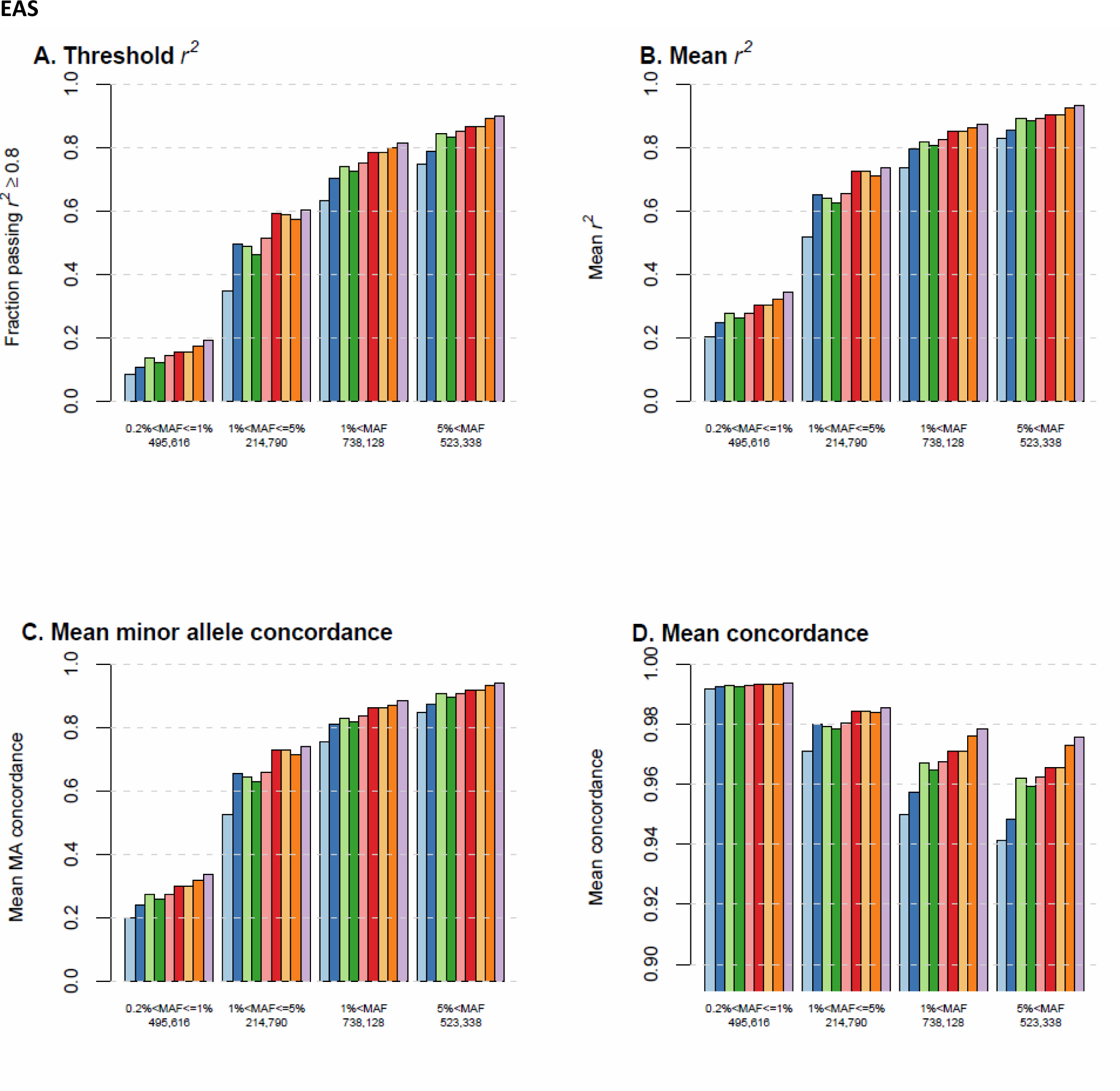

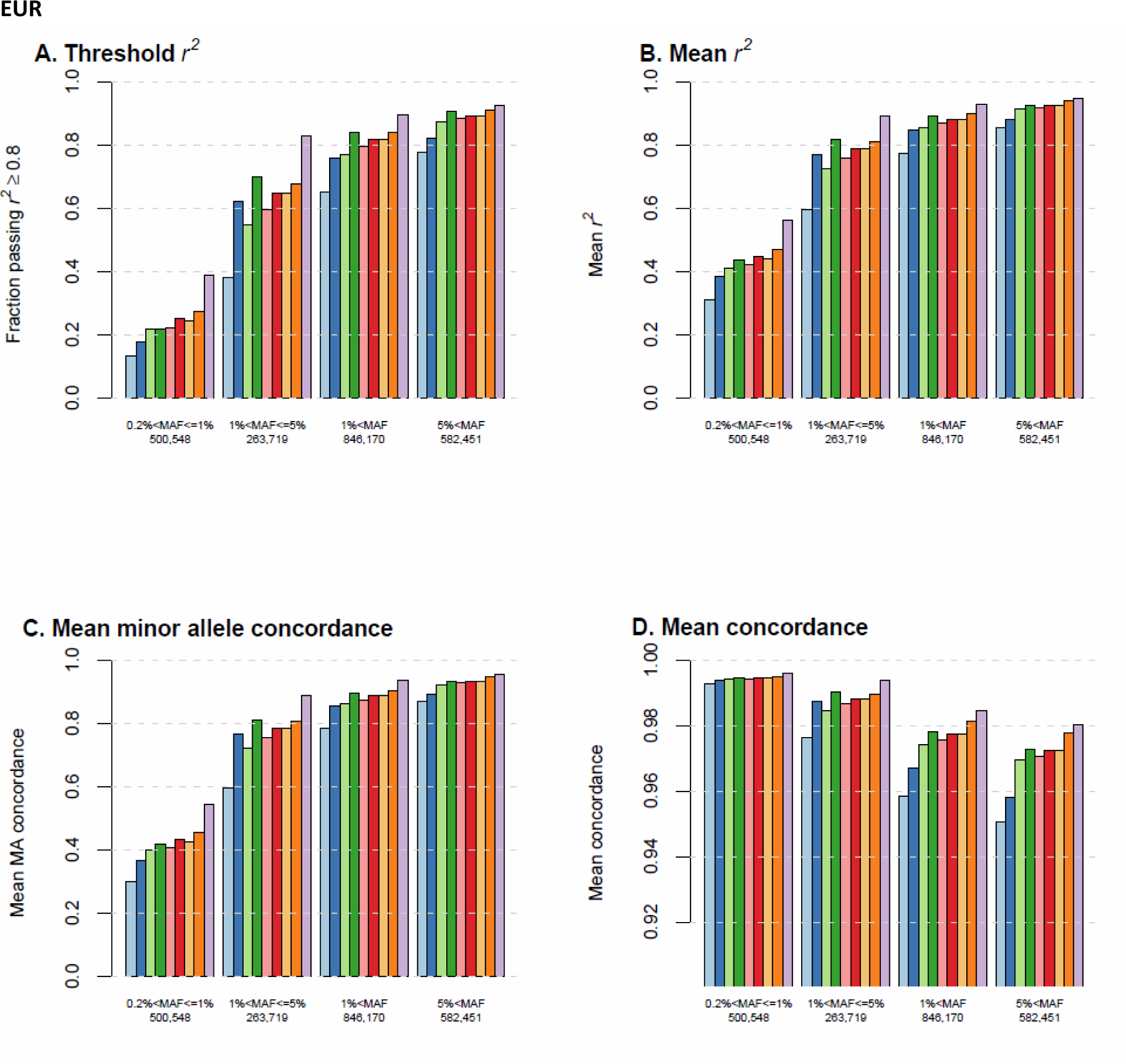

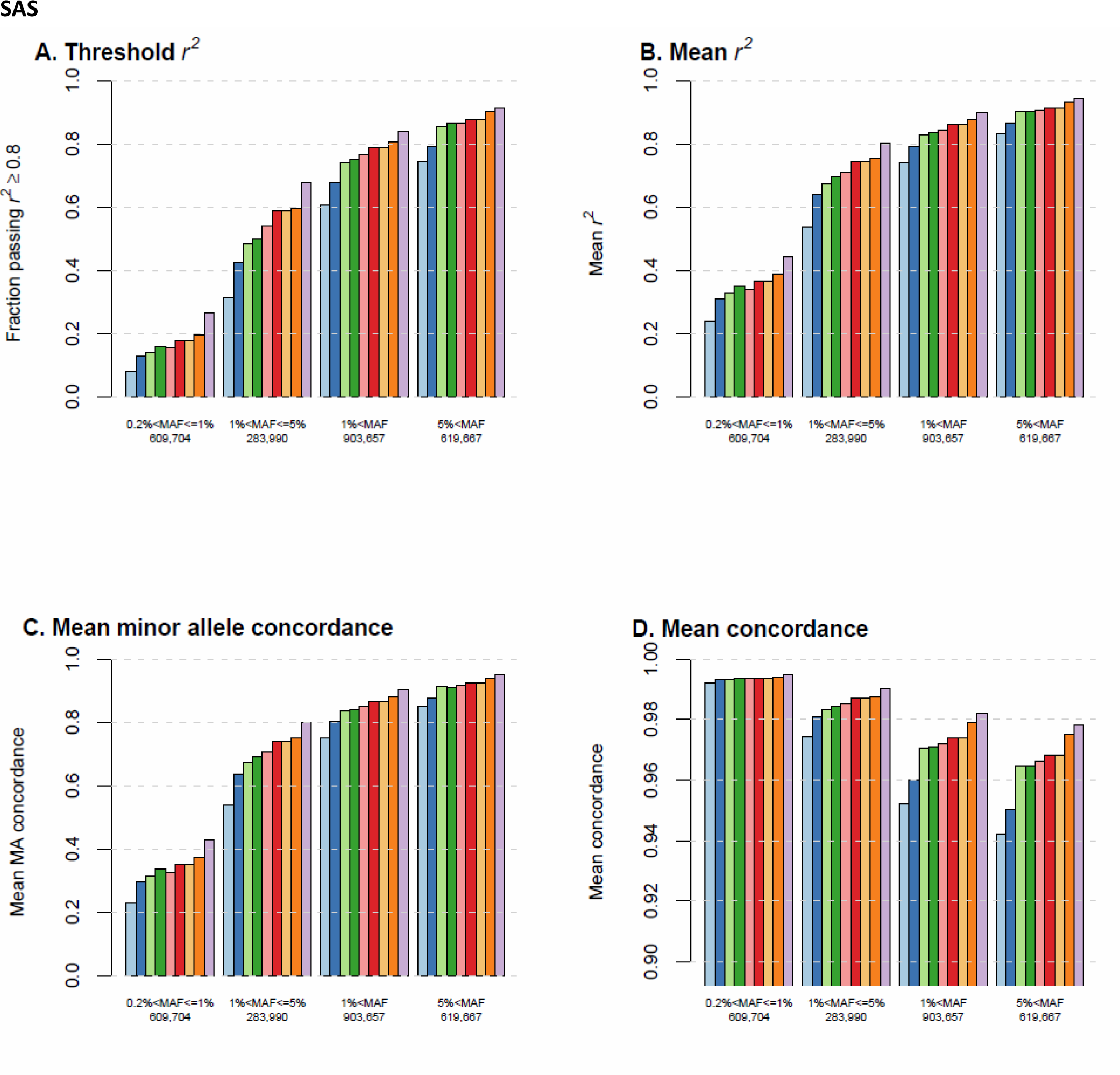

